# REGENERATION VIA SOMATIC EMBRYOGENESIS FROM SEED EXPLANT OF MEDICINAL PLANT *SOLANUM VIRGINIANUM* (L.)

**DOI:** 10.1101/2022.07.22.500937

**Authors:** Dhanashree S. Patil, Swaroopa A. Patil

## Abstract

*Solanum virginianum* (L.) belonging to family Solanaceae is an important medicinal plant, used in traditional ayurvedic medicines. It is one of the ten constituents of “Dashmoola”. Present work deals with the development of efficient procedure for inducing somatic embryogenesis followed by regeneration of *Solanum virginianum* (L.). Initially, seeds of *S. virginianum* were used as explant and inoculated on ½ MS basal medium. The stem explant from in vitro germinated seeds was used for callus induction. MS medium supplemented with BAP (1mg/L) was used for callus induction. Callus proliferation started when initiated callus was subcultured on different concentrations and combination of BAP, IAA, IBA, 2,4-D and NAA. Where highest embryogenesis took place in BAP (2mg/l)+IAA(5mg/l) combination. The somatic embryos were transferred onto MS basal medium for maturation. These embryos matured and were converted into complete plantlets with well developed shoot and root system.

## 1. Introduction

Medicinal plants are wealth of humankind. India has largest biodiversity and has high knowledge of rich ancient traditional system of medicine (Ayurveda, Siddha, Unani, Amchi and local health traditions) which provides a strong base for the utilization. *Solanum virginianum* (L.), which belongs to the family Solanaceae, is an important medicinal plant species in ayurveda and folklore medicine since time immemorial (Rita and Animesh, 2011). Although the plant is very important traditionally but lack of experimental research is a hindrance for its exploration in modern system of medicine. *Solanum virginianum* (L) is commonly known as yellow berried night shade. *Solanum surattense* Burm. (F) and *Solanum xanthocarpum* Schrad. and Wendl. are synonyms of *Solanum virginianum* L. (Sharma *et al*., 2010). It is frequently distributed in Asia including India, China, Nepal, Pakistan, Sri lanka, Bangladesh, Afghanistan, Thialand etc. In India it is recorded in tropical, subtropical, and all the four geographical regions mostly in dry places. Morphologically *Solanum virginianum* (L.) is a prickly diffuse bright green perennial herb which is erect or creeping, sometimes woody at base which is 1.2m in height, copiously armed with sturdy, needle like, broad based prickles which are straight and compressed, glabrous and shining often 1-3cm long. Leaves are unequal paired, leaf blade ovate or elliptic, subpinnatified oblong, stellately hairy on both the sides, armed on the midrib, the nerves with long yellow sharp prickles, flowers are blue-purple in cymes and sometimes reduced in solitary, fruit types are gobular berry 1.25-2cm in diameter, The berries are green and white strips when unripe and yellow when gets ripened. Branches are numerous and young ones are covered with dense, stellate and tomentose hairs. (Rane *et. al*., 2014).

*S. virginianum* (L.) is the plant that has many medicinal properties. It is one of the ten constituents of “Dashmoola”. Ayurvedic physicians commonly use the drugs of Dashmoola in their private practices. The tribal people use the drugs of Dashmoola group in their common ailments. *S. virginianum* is a chemically valuable source of alkaloids, sterols, saponins, flavonoids and their glycosides (Okram and Thokchom, 2010). It also contains carbohydrates, fatty acids and amino acids (Gnana *et al*., 2013). Heble *et al*., (1968) chemically isolated, crystallized, diosgenin and beta cytosterol are found in *S. virginianum* (L.). Triterpene like Lupeol has been reported (Heble *et al*., 1971). Apart from this it contains coumarins, scopolin, scopoletin, esculin and esculetin (Tupkari *et. al*., 1972), alkaloids, flavoinoids and saponin and tolerable levels of heavy metals like Cu, Fe, Pb, Cd and Zn (Hussain *et al*., 2010). Bioactive steroidal glycoalkaloid khasianine in addition to solanine and solasomargine has been reported in *S. virginianum* (L.) (Shankar *et. al*., 2011). Solasodine serves as an important intermediate in the synthesis of steroidal hormone (Butcher 1977) and is an alternative to diosgenins as a precursor in synthesis of steroidal hormones like progesterone and cortisone (Marck, 1989; Galanas *et al*., 1984). Apigenin, another secondary metabolite from *S. virginianum* has antiallergic property while diosgenin exhibit anti – inflammatory effects (Singh *et al*., 2010).) *Solanum virginianum* has larvicidal activity against snails and mosquito larvae (Tanusak Changbanjong *et al*., 2010). It is traditionally used for treating tuberculosis (TB), fever, bronchial asthma, lung diseases and kidney disorders. (Govindan *et al*., 2004). Krayer and Briggs (1950) reported the antiaccelerator cardiac action of solasodine and some of its derivatives. The plant possesses antiurolthiatic and natriuretic activities (Patel *et al*., 2010). A decoction of the fruits of the plant is used for treatment of diabetes (Nadkarni, 1954). *Solanum virginianum* (L) herb is useful in cough, chest pain, against vomiting, hair fall, leprosy, itching scabies, skin diseases and cardiac diseases associated with edema (Kumar *et al*., 2010). Roots decoction is used as fabrige, effective diuretic and expectorant (Ghani, 1998). Root paste is utilized by the Mukundara tribals of Rajasthan for the treatment of hernia as well as in flatulence and constipation. Stem, flower and fruits are prescribed for relief in burning sensation in the feet. Leaves are applied locally to relieve body or muscle pains, while its juice mixed with black pepper is advised for rheumatism (Nadkarni, 1954). Fruit juice is useful in sore throats and rheumatism. A decoction of the fruits of the plant is used by tribal and rural people of Orissa for the treatment of diabetes (Nadkarni, 1954). Smoking the seeds of the dried *S. virginanum* in a biri wrap is used to relieve toothache and tooth decay in Indian folk medicine. Fruit, flowers and stem possesses carminative, anthelmic and bitter properties, root is expectorant and used in the treatment of toothache. The leaves are used externally to relieve pain. Propagation of *S. surattense*, takes place only by seeds but it may face some difficulties such as low rate of germination under normal conditions, loosing of viability on storage, the progenies produced may not be true to type due to cross pollination (Yadav *et al*., 2010). Due to it’s over exploitation for high medicinal values this plant is becoming endangered (Khan and Frost, 2001), hence there is a need for ex-situ conservation through tissue culture technique (Sinha *et al*., 1979).

Globally, plant tissue culture (PTC) have been recognized for its valuable and beneficial role in the production and conservation of plant based resources (Vasil, 2008). PTC can be achieved with the help of different explants such as meristem culture, anther culture, ovule culture, endosperm culture, nucellus culture, embryo culture etc. Somatic embryogenesis is a developmental process of producing normal embryos from undifferentiated somatic cells. Somatic embryogenesis is the maximum expression of cell totipotency in plant cells. In short, totipotency is the ability of a plant cell to undergo a series of complex metabolic and morphological coordinated steps to produce a complete and normal plant or sporophyte without the participation of the sexual processes. Thus, somatic embryogenesis is the developmental process by which theoretically any somatic cell develops into a zygotic like structure that finally forms a plant (Rao, 1996; Jimenez, 2005). Somatic embryogenesis is significantly important for *in vitro* mass propagation, germplasm conservation and genetic improvement of higher plants.

Very limited reports are available on the *In vitro* regeneration of *Solanum virginianum*. Using nodal and shoot tip segments, micropropagation technique has been established in *S. virginianum* (Ramaswamy *et al*., 2004). Direct shoot organogenesis from shoot tip and leaf segment was achieved by Pawar *et al*. (2002). Somatic embryogenesis was successfully achieved from cotyledon and leaf explants in *S. surattense* (Ramaswamy *et al*., 2005). Callus induction and shoot organogenesis from floral bud was studied (Prasad *et al*., 1998). The present study is focused on the effect of growth regulators that enhance somatic embryogenesis followed by shoot initiation in *S. virginianum* for mass propagation.

Somatic embryogenesis is an alternative and efficient method for plant propagation. The plants regenerated via somatic embryogenesis are of single cell origin (Rao, 1996), true to type and are produced in large number within short period of time.

## 2. Materials and Methods

### 2.1 Collection of plant material

Morphologically healthy fruits of *Solanum virginianum* (L.) were collected from Haripur, Sangli (16^0^ 50’ 52.2” N 74^0^ 32’ 05.8” E) (Maharashtra, India). The plant sample was submitted to SUK acronym (DSP 01). The raised plantlets are maintained at botanical garden of Department of Botany, Shivaji University, Kolhapur.

### 2.2 Selection of explant and inoculation

Healthy and disease free seeds from the maintaind germplasm were selected as explant. These were used for inoculation.

#### 2.2.1 *In-vitro* seed germination

##### 2.2.1.1 Sterilization

All the fruits were washed with detergent followed by tap water twice in laboratory and surface sterilized with 0.1% HgCl2 for 4 minutes, followed by 3 washes of sterilized distilled water for 1 minute each and then treated with 70% ethanol for 30 seconds followed by 3 washes of sterilized distilled water for 1 minute each. Surface sterilization was carried out in laminar air flow. Then all the fruits were blotted dry. The fruits were then cut and seeds were dissected out. The seeds were inoculated on to ½ strength MS medium containing 3% sucrose and 0.8% Agar agar aseptically.

##### 2.2.1.2 Induction of callus

After germination, the stem from the germinated seedling was used as explant and it was cultured on MS medium supplemented with BAP (1mg/l).

##### 2.2.1.3 Induction of Somatic Embryogenesis

For induction of somatic embryogenesis the callus formed was transferred on MS medium supplemented with various plant growth regulators either singly or in combination. Once the nodular embryos were formed, for further development of embryos they were transferred on to the same hormone concentration or hormone free MS medium.

### 2.3 Culture condition

After inoculation culture tubes were transferred to culture room under a 16/8hr (light/dark) photoperiod, temperature 25±2°C and maximum humidity (80%) was adjusted with air conditioner.

### 2.4 Statistical analysis

The observations were recorded after 9 weeks and statistical analysis was done using Microsoft excel statistical methods. The values were represented as mean ±SE. Each experiment had 20 replicates and was repeated three times.

## 3. Results and discussion

The important step in plant tissue culture was surface sterilization of explants. Minimum contamination and higher survival rate was achieved in 0.1% of HgCl2. However the optimum time period of exposure of explants to HgCl_2_ was 3 minutes. Surface sterilized explant was inoculated on ½ strength MS medium. Within 15 to 20 days seeds started germinating. After germination of seeds the elongated shoots were inoculated on different concentrations of BAP (0.5mg/l, 0.75mg/l, 1mg/l) for multiplication. Maximum growth was achieved on 0.75mg/l BAP. BAP has been more effective for shoot regeneration than Kn for shoot multiplication of *S. virginianum* (Sundar *et al*., 2011). Ramar and Nandagopalan (2011) confirmed that *Solanum surattense* has low amount of endogenous hormones hence it requires high amount of exogenous hormones for plant regeneration. The stems from germinated seedlings were inoculated on 1mg/l BAP which resulted in callus induction. The explants were slightly swollen and formed protuberance along whole length of the explants, which upon continuous culture gave rise to pale greenish callus within 3 to 4 weeks. The induced callus was subcultured on various concentrations and combinations of BAP, IAA, IBA, 2,4-D and NAA used singly or in combination (Fig. 1, Table 1). Proliferative callus was converted into high nodular embryogenic callus in all the combinations tried. When combinations like BAP+IAA and BAP+IBA were further used for subculturing, proliferative callus was seen to rapidly convert into embryogenic callus. In medium with single hormone (NAA/2,4-D) it was seen that, proliferative callus subcultured on hormone free medium (devoid of NAA/2,4-D) showed somatic embryogenesis. Thus to initiate proliferative calli BAP, IAA, IBA, NAA and 2,4-D were used alone or in combination. For achieving somatic embryogenesis BAP+IAA, BAP+IBA combinations were used for at the time of subculturing. While in case of single hormone used the subculturing medium was devoid of any plant growth regulator. All the combinations of growth regulators were more or less responsible for callus proliferation. The embryogenic callus was composed of small spherical or globular nodules. Callus formation process needed 4-6 weeks for best proliferative callus. The embryogenesis was seen in the callus in the 7^th^ week in some combinations which extended to the 9^th^ week where somatic embryogenesis was observed in all the hormones tried. Dark conditions favored induction of somatic embryogenesis.

**Fig. 1.**
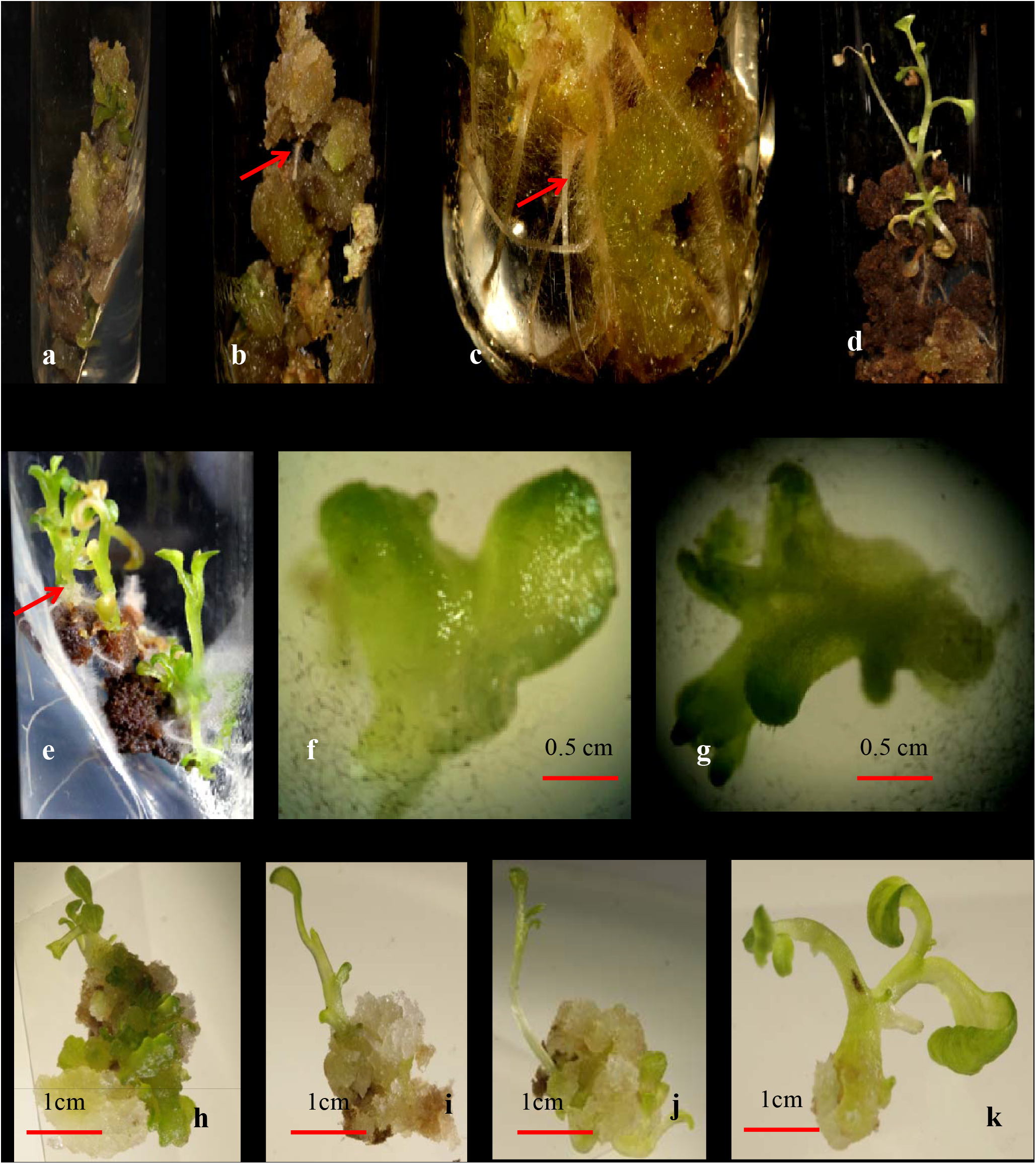
a-k Somatic embryogenesis in Solanum virginianum (L.) a. Embryogenenic callus, b. Induction of polar tip of embryo, c. Root development form the polar tip of embryo, d. Shoot initiation from the opposite pole of cell, e. Embryo growing into a well developed plantlet, f. Heart shaped somatic embryo,. g. Torpedo stage of embryogenesis, h.,i. and j. Developing shoot tips, k.Distinct shoot tip dissected out from callus mass.

**Table 1.**
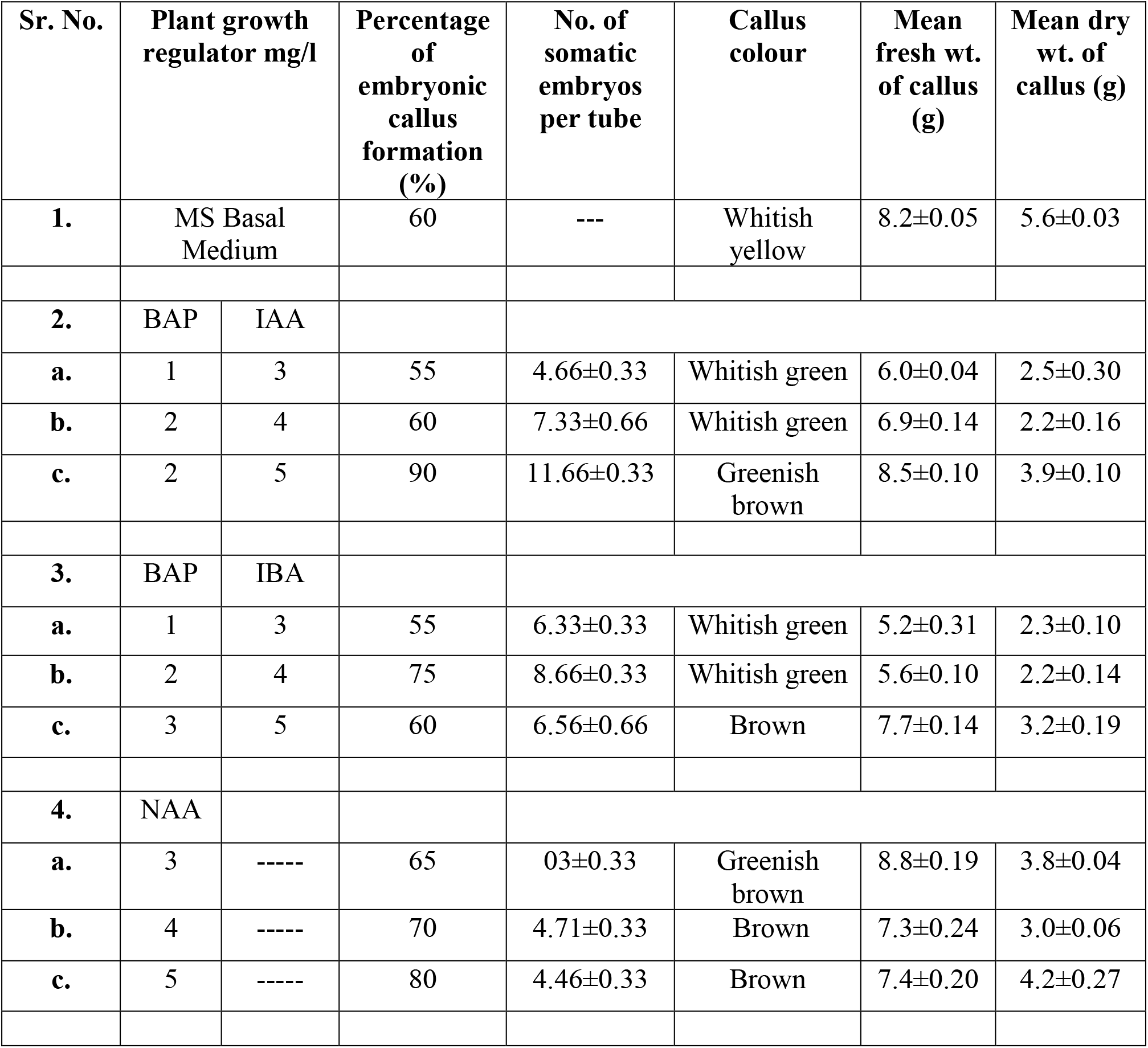
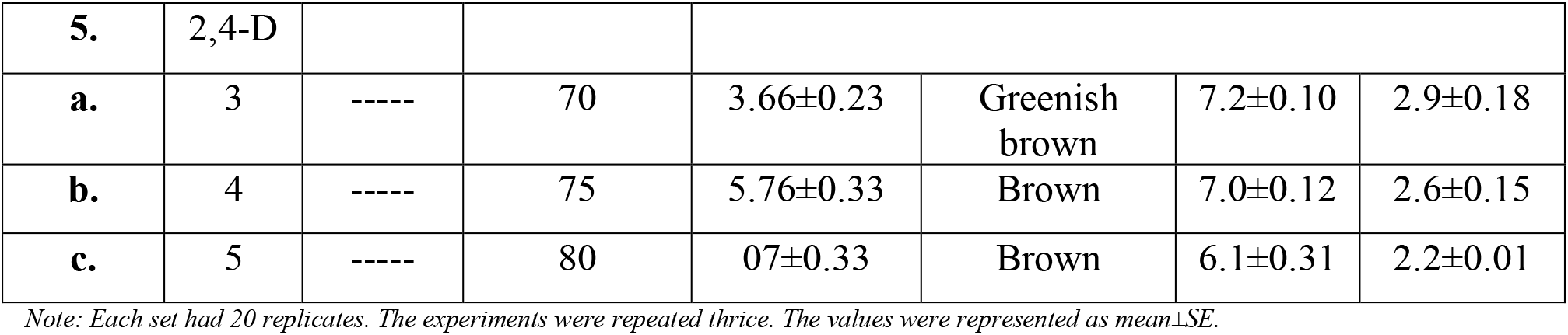
Effect of different concentrations and combinations of Growth Regulators on callus of *Solanum virginianum*.

In all combinations of BAP+IAA studied, maximum % response for embryogenesis (90%) was recorded in 2mg/l BAP in combination with 5mg/l IAA. Cytokinin, such as BAP is reported to inhibit embryogenic potential in carrot cultures but a combination of BAP with IAA was found to induce embryogeneses in *Podophyllum hexandrum* (Dantu and Tomar, 2010). Combinations of BAP+IAA for somatic embryogenesis in present findings are in accordance to this report.

Embryogenic callus induction was also tried with 2,4-D and NAA alone, whitish callus was observed. Maximum growth was recorded on 5mg/l for both. Ramaswamy *et al*. (2005) showed that, leaf explants of *Solanum surattense* showed maximum percentage of somatic embryogenesis and high frequency of somatic embryo induction when MS medium supplemented with 4mg/l NAA and 0.5mg/l BAP. Somatic embryogenesis was reported on the medium containing NAA alone in *Solanum melongena* (Sharma and Rajam, 1995).

Less concentration of growth regulators were showing poor growth of callus. Initial dependence of callus for growth hormone to initiate embryogenesis was reported in number of species but it was found necessary to withdraw the growth hormone from the medium for subsequent growth of somatic embryos. Ghatge *et al*. (2008) showed that in the presence of 2,4-D initial embryogenic competence of callus was achieved, but it was found necessary to withdraw 2,4-D from the medium for the further development of somatic embryos (Zimmerman, 1993), but according to our observations growth hormones that is BAP+IAA and BAP+IBA are required for the somatic embryogenesis when combinations of hormones were used. Whereas withdrawal of the growth hormones NAA as well as 2,4-D leads to somatic embryogenesis in case of single hormone used.

In the present study, morphological observations showed the heart-shaped somatic embryos further progressed towards late-heart-shaped or torpedo-shaped embryos as they attained well-developed and mature cotyledons. Similar study was carried in *Solanum tuberosum*, where two cotyledonary stages were observed onto the medium supplemented with 5μM 2,4-D. (Sharma and Millam, 2004).

A balance in endogenous and exogenous auxin is very important for somatic embryogenesis. In citrus culture it was observed that, at very low concentration of auxin, cultured for continuous two years, failed to give somatic embryogenesis but when the same culture supplemented with auxin inhibitor, somatic embryogenesis was achieved (Kochba and Spiegel-Roy, 1977). In carrot cultures it was seen that, auxin containing cultures produces more ethylene than auxin free cultures. High ethylene content enhances more activity of cellulase or pectinase or both which causes breakdown of clumps before polarity is established in the proembryos for further development. Thus 2,4-D supports tissue multiplication but opposes maturity of the embryos (Dantu and Tomar, 2010). Cytokinins had significant effects on somatic embryogenesis and high concentration (1mg/l) was more effective. Similar results were obtained by Singh and Chaturvedi (2009) who reported that the effects of cytokinins on somatic embryogenesis were significant and mandatory for the differentiation of somatic embryos. In conclusion, the present study showed that, callus induction and SE were strongly affected by plant growth regulators and culture conditions. The medium containing both auxin and cytokinin at any concentration was capable of inducing the formation of somatic embryogenesis but differences were observed based on the type and concentration of plant growth regulators and light conditions (Nuray *et al*., 2009).

## 4. Conclusion

It was observed that, dark conditions favoured induction of somatic embryos. The time taken for the induction of somatic embryos was between 4 to 6 weeks. The development of somatic embryos was seen in all the cultures with varied responses. The present study revealed that, different concentrations of BAP with IAA, IBA induces embryogenic callus followed by somatic embryogenesis and 2,4-D and NAA alone induces embryogenic callus later withdrawal of the same results in somatic embryogenesis. The protocol developed is efficient and can be exploited for large scale multiplication of this medicinal plant.

## Supporting information

Abstract

Author Information

