## Supplementary material for "REGENERATION VIA SOMATIC EMBRYOGENESIS FROM SEED EXPLANT OF MEDICINAL PLANT *SOLANUM VIRGINIANUM* (L.)": Author Information

**Dhanashree S. Patil**

**Research scholar**

**Department of Botany**

**Shivaji University Kolhapur**

**416004.**

**Mob. No. 7219835288**

**Dr. Swaroopa A. Patil***

**Assistant Professor**

**Department of Botany**

**Shivaji University Kolhapur**

**416004.**

**Mob. No. 8830778706**

***Corresponding author**

**Running title:** Somatic embryogenesis in *Solanum virginianum* (L.).
