## Supplementary material for "REGENERATION VIA SOMATIC EMBRYOGENESIS FROM SEED EXPLANT OF MEDICINAL PLANT *SOLANUM VIRGINIANUM* (L.)": Table 1

**Table 1 Effect of different concentrations and combinations of Growth Regulators on callus of *Solanum virginianum***

| **Sr. No.** | **Plant growth regulator mg/l** | | **Percentage of embryonic callus formation (%)** | **No. of somatic embryos per tube** | **Callus colour** | **Mean fresh wt. of callus**  **(g)** | **Mean dry wt. of callus (g)** |
| --- | --- | --- | --- | --- | --- | --- | --- |
| **1.** | MS Basal Medium | | 60 | --- | Whitish yellow | 8.2±0.05 | 5.6±0.03 |
| **2.** | BAP | IAA |  |  | | | |
| **a.** | 1 | 3 | 55 | 4.66±0.33 | Whitish green | 6.0±0.04 | 2.5±0.30 |
| **b.** | 2 | 4 | 60 | 7.33±0.66 | Whitish green | 6.9±0.14 | 2.2±0.16 |
| **c.** | 2 | 5 | 90 | 11.66±0.33 | Greenish brown | 8.5±0.10 | 3.9±0.10 |
| **3.** | BAP | IBA |  |  | | | |
| **a.** | 1 | 3 | 55 | 6.33±0.33 | Whitish green | 5.2±0.31 | 2.3±0.10 |
| **b.** | 2 | 4 | 75 | 8.66±0.33 | Whitish green | 5.6±0.10 | 2.2±0.14 |
| **c.** | 3 | 5 | 60 | 6.56±0.66 | Brown | 7.7±0.14 | 3.2±0.19 |
| **4.** | NAA |  |  |  | | | |
| **a.** | 3 | ----- | 65 | 03±0.33 | Greenish brown | 8.8±0.19 | 3.8±0.04 |
| **b.** | 4 | ----- | 70 | 4.71±0.33 | Brown | 7.3±0.24 | 3.0±0.06 |
| **c.** | 5 | ----- | 80 | 4.46±0.33 | Brown | 7.4±0.20 | 4.2±0.27 |
| **5.** | 2,4-D |  |  |  | | | |
| **a.** | 3 | ----- | 70 | 3.66±0.23 | Greenish brown | 7.2±0.10 | 2.9±0.18 |
| **b.** | 4 | ----- | 75 | 5.76±0.33 | Brown | 7.0±0.12 | 2.6±0.15 |
| **c.** | 5 | ----- | 80 | 07±0.33 | Brown | 6.1±0.31 | 2.2±0.01 |

*Note: Each set had 20 replicates. The experiments were repeated thrice. The values were represented as mean±SE.*
